# Changes in the relative abundance of two *Saccharomyces* species from oak forests to wine fermentations

**DOI:** 10.1101/034157

**Authors:** Sofia Dashko, Ping Liu, Helena Volk, Lorena Butinar, Jure Piškur, Justin C. Fay

## Abstract

*Saccharomyces cerevisiae* and its sibling species *S. paradoxus* are known to inhabit temperate arboreal habitats across the globe. Despite their sympatric distribution in the wild, *S. cerevisiae* is predominantly associated with human fermentations. The apparent ecological differentiation of these species is particularly striking in Europe where *S. paradoxus* is abundant in forests and *S. cerevisiae* is abundant in vineyards. However, ecological differences may be confounded with geographic differences in species abundance. To compare the distribution and abundance of these two species we isolated *Saccharomyces* strains from over 1,200 samples taken from vineyard and forest habitats in Slovenia. We isolated numerous strains of *S. cerevisiae* and *S. paradoxus* as well as small number of *S. kudriavzevii* strains from both vineyard and forest environments. We find *S. cerevisiae* less abundant than *S. paradoxus* on oak trees within and outside the vineyard, but more abundant on grapevines and associated substrates. Analysis of the uncultured microbiome shows that both *S. cerevisiae* and *S. paradoxus* are rare species in soil and bark samples, but can be much more common in grape must. In contrast to *S. paradoxus*, European strains of *S. cerevisiae* have acquired multiple traits thought to be important for life in the vineyard and dominance of wine fermentations. We conclude that *S. cerevisiae* and *S. paradoxus* currently share both vineyard and non-vineyard habitats in Slovenia and we discuss factors relevant to their global distribution and relative abundance.

## Introduction

The ability to ferment sugar in the presence of oxygen originated around the time of a whole genome duplication and is shared by many yeast species (Hagman et al., 2013). Among these yeasts, *Saccharomyces* species are distinguished in being present and intentionally used by humans for the production of alcoholic beverages. While strains of *S. cerevisiae* are the most widely used, other *Saccharomyces* species and their hybrids are involved in various types of fermentations. *S. cerevisiae* x *S. eubayanus* hybrids are used for lager production (Libkind et al., 2011; Nakao et al., 2009), *S. cerevisiae* x *S. kudriavzevii* and *S. cerevisiae* x *S. uvarum* hybrids are used for low temperature wine production (Belloch et al., 2009; Borneman et al., 2012; Bradbury et al., 2006; Erny et al., 2012; Le Jeune et al., 2007; Lopandic et al., 2007; Oliva et al., 2006), and *S. uvarum* is used to ferment apple cider and wine at low temperatures (Naumov et al., 2000, 2001; Rainieri et al., 1999). One of the clearest differences among these species and one taken advantage of for certain types of fermentations is their thermal growth profile; *S. cerevisiae* and *S. paradoxus* are thermophilic and *S. kudriavzevii* and *S. uvarum* are cryophilic (Gonçalves et al., 2011; Salvadó et al., 2011). However, other aspects of the ecology and evolution of these species might also be relevant to the origin of industrial yeast strains and the predominant use of *S. cerevisiae*.

Outside human ferments, the *Saccharomyces* species have primarily been isolated from arboreal habitats. Originally noted to be associated with sap seeping from slim fluxes (Naumov et al., 1998), these yeast species have now been consistently isolated from bark, leaves and surrounding soil of *Quercus* (Oak) and other tree species. Both *S. cerevisiae* and *S. paradoxus* are widely distributed and have been isolated from temperate forests in North America (Charron et al., 2014; Hyma and Fay, 2013; Sniegowski et al., 2002; Sylvester et al., 2015), Europe (Almeida et al., 2015; Bozdag and Greig, 2014; Johnson et al., 2004; Koufopanou et al., 2006; Legras et al., 2014; Naumov, 2013; Sampaio and Gonçalves, 2008), Asia (Almeida et al., 2015; Naumov et al., 1997; Wang et al., 2012) and Oceania (Zhang et al., 2010), often sympatrically. *S. uvarum* and *S. eubayanus* have also been found to be widely distributed (Almeida et al., 2014). However, *S. kudriavzevii*, *S. arboricola* and *S. mikatae* have thus far only been found in restricted geographic regions (Naumov et al., 2013). Currently, it is unknown whether arboreal habitats are a primary habitat or just one of many environments able to sustain populations of these species (Goddard and Greig, 2015).

Vineyards are likely an important interface between wild yeast populations and those used for wine fermentations (Hyma and Fay, 2013). Grapes periodically provide an abundant source of sugar, attract a high density of potential insect vectors, and generate exceptionally high concentrations of yeast by the end of fermentation. Furthermore, wine must is not sterilized prior to fermentations and the skins, stems and microbial sediments from fermentation are typically discarded back into the vineyard. Thus, before the now common practice of inoculating wine must, there was ample opportunity for both inter-and intraspecific competition within vineyard environments. Indeed, commercial wine yeast is found dispersed throughout vineyards in France (Valero et al., 2005), and European “wine” strains and North American “wild” strains of *S. cerevisiae* are both present on grapes and oak trees in vineyards in North America, while only North American “wild” strains are found in arboreal habitats outside of vineyards (Hyma and Fay, 2013). Mixing of various *S. cerevisiae* populations also occurs in Italy, facilitated by wasps (Stefanini et al., 2012). Finally, the above mentioned hybrids of *Saccharomyces* species have thus far only been isolated from vineyard and brewing environments.

The historical acquisition of *S. cerevisiae* but not *S. paradoxus* into human-associated fermentative environments is particularly perplexing given they are both strong fermenters and widely distributed. For example, *S. paradoxus* has only been reported as a significant contributor to wine fermentations in Croatia (Redzepović et al., 2002). Furthermore, many of the growth characteristics that give *S. cerevisiae* a competitive advantage during wine fermentations are shared with *S. paradoxus* and the two species are equally competitive in high sugar environments such as grape juice (Williams et al., 2015). Consistent with these observations, both *S. cerevisiae* and *S. paradoxus* have been isolated from vineyards in North America (Hyma and Fay, 2013).

In Europe, however, there appears to be ecological differentiation between wine strains of *S. cerevisiae* and wild populations of *S. paradoxus*. Historically, *S. paradoxus* was isolated from arboreal habitats while *S. cerevisiae* was isolated from vineyards (Naumov, 2013), which lead to the reasonable proposition that *S. cerevisiae* is a domesticated species (Martini, 1993; Mortimer, 2000). While absent from northern European arboreal habitats (Johnson et al., 2004; Koufopanou et al., 2006; Sampaio and Gonçalves, 2008), *S. cerevisiae* has now been isolated from multiple Mediterranean oak trees (Almeida et al., 2015; Sampaio and Gonçalves, 2008) and may constitute a wild stock from which European wine strains were derived. In contrast, *S. paradoxus* has been isolated from arboreal habitats throughout Europe (Boynton and Greig, 2014; Glushakova et al., 2007; Naumov, 1996, 2013; Naumov et al., 1992; Sampaio and Gonçalves, 2008). One caveat, however, is that concurrent sampling of vineyard and arboreal habitats within the same region is needed to tease apart geographic and habitat effects on the abundance and distribution of these species.

In this study, we examine the abundance of *S. cerevisiae* and *S. paradoxus* across sympatric ecological environments and fine-scale geographic locations in Slovenia. Our sampling strategy was arboreal sources, including bark and soil from oak trees, within and outside of vineyards compared to wine must, soil and berries from grapevines within vineyards. Using enrichments we find both species present within and outside the vineyard, and analyze their abundance in arboreal-and grape-associated habitats. We also quantify species abundance using enrichment free microbial profiling of bark, soil and wine must before and during fermentation. By quantifying phenotypes relevant to life in the vineyard we provide an explanation for why *S. paradoxus* is rare or absent in autochthonous wine fermentations despite its presence in the vineyard.

## Materials and Methods

### Sample collection and strain isolation

Samples were obtained from seven vineyards and four forest sites in Slovenia (Table S1). The majority of samples were soil, bark and berries from grapevines and soil and bark from oak trees (*Quercus robur*, *Q. petrea*, *Q. ilex*, *Q. pubescens* and *Q. cerris*). A small number of samples were from insects, fruits, cellar swabs and wine must. Oak samples were obtained by prying off bark at the base of the tree and sampling soil at the base of the tree. Samples were obtained between July of 2013 and April of 2014. Three of the forest locations were transects from the edge of Vipava Valley leading up into the surrounding mountains and these forested areas began immediately adjacent to vineyards. One forest location was on a hill above the town of Vipava surrounding an abandoned castle.

For each sample, approximately 5–25 cm^3^ of substrate was placed into a sterile falcon tube using ethanol sterilized forceps or scalpels. Twenty-five ml of enrichment medium (1% yeast extract, 2% peptone, 10% glucose and 5% of ethanol, pH 5.3) was added to each sample (Hyma and Fay, 2013; Mortimer and Polsinelli, 1999). After four to ten days of fermentation at room temperature, approximately 20–25°C, 2 µl of well mixed enrichment medium was spread on Petri dishes and incubated for 2–4 days. Bacteria-like colonies were excluded by testing for growth on chloramphenicol (100 mg/L). A single yeast colony was isolated from each enrichment and place into 3 ml of liquid YPD and incubated with 200 rpm shaking overnight. For 13 enrichments we isolated two colonies from the same enrichment corresponding to different morphology. Only one of the two isolates was used in the analysis.

A subset of 518 samples collected in October were enriched at both room temperature and 37°C. These samples were derived by thoroughly mixing each sample with enrichment media, then pouring off 10 ml of the enrichment into a sterile, 15 ml tube and incubated at high temperature. The high temperature enrichments were subsequently treated the same as those at room temperature and single colonies were obtained from both.

### Species identification

Isolates were screened for *Saccharomyces* species by PCR and restriction digests as in Hyma and Fay (2013). Briefly, total DNA was extracted from yeast using lyticase and glass beads. A multiplex PCR assay was used to distinguish *Saccharomyces* and non-*Saccharomyces* species (Nardi et al., 2006). *Saccharomyces* isolates were further distinguished using restriction digests of the ITS PCR products (McCullough et al., 1998). Identification failed for 56 (6%) isolates, either because of PCR failure or digests with unexpected band sizes.

### Sampling analyses

For each species, the frequency of isolation from all oak-and grapevine-associated samples was fit to a logistic model with terms for source (oak, grapevine), location type (vineyard, non-vineyard), location (11 sampling sites) and month of isolation (July, September, October, April). Significant terms were identified by dropping single terms and comparing models using a likelihood ratio test.

### Microbiome analysis

Microbiome samples were collected after harvest from vineyards and wine must. From five vineyards we obtained twenty samples from oak bark and soil and grapevine soil and twenty must samples from uninoculated fermentations. Nine of the must samples were from pressed grapes or pomace within a day of harvest, the remainder were from within the first week of fermentation. Temporal samples were taken from fermenting must from two vineyards, Carga and Burja (previously part of the Sutor estate). From Burja, samples were taken from Malvazija pomace at harvarst (20.2 °Brix, pH 3.37, total titratable acidity 7.1 g/L, supplemented with ammonium bisulfite at 0.1 g/L) and at seven subsequent points over 18 days of fermentation in the cellar. The same must was also brought to the lab and 800 ml of must was fermented in flasks in triplicate during which we obtained 8 samples over 14 days. On the eleventh day of fermentation, must in the lab and in the cellar was pressed to remove the skins and seeds and the remaining juice continued to ferment. From Carga, juice from pressed Tokaj grapes 17.9 °Brix, pH 3.21, total titratable acidity 8.1 g/L, supplemented with potassium metabisulfite at 0.1 g/kg) was brought to the lab and 800 ml of must was fermented in flasks in triplicate during which we obtained 10 samples over 17 days.

For soil samples, DNA was extracted from 150 mg of soil using ZR Soil Microbe DNA extraction kit (Zymo Research, CA, USA). For bark, berry and juice/pomace samples, samples were immersed and shaken in water, solid material removed, and DNA was extracted from the pellet after centrifugation using either the ZR Soil Microbe kit (bark) or a Qiagen Plant DNA kit (Hilden, Germany). ITS1 was amplified using BITS1 and B58S3 primers (Bokulich and Mills, 2013). For the BITS1 primer we included an 8 bp barcode followed by a linker (CT) at the 5’ end (Table S2) in order to multiplex the samples. Illumina sequencing adaptors were added via a second round of PCR and these included a 9 bp index for further multiplexing. Amplicons were purified, quantified and pooled then sequenced using an Illumina MiSeq with single-end 250 bp reads.

Barcodes and index were identified and removed using custom perl scripts allowing 1 mismatch in each. Adaptors and low quality sequences were trimmed using ea-utils (v1.04.676 https://code.google.com/archive/p/ea-utils/) using a window size of 3 and a quality threshold of 20. Sequences less than 100 bp were removed. Sequences were aligned by blastall (v2.2.26 (Camacho et al., 2009)) using a cutoff of 1e-20 to 287,101 sequences in the UNITE+INSD database (4/7/2014 (Kõljalg et al., 2013)) after removing sequences from uncultured fungi. For classification into taxonomic groups all top hits were used. For species classification, the top hit for each sequence was retained when greater than 97% identity, resulting in the retention of 75% of all sequences. To eliminate rare and potentially spurious hits, species representatives were only kept if two or more samples had more than 10 hits each to that representative. This eliminated 3,091 out of 3,935 species representatives. After these filters, the median number of hits per sample was 46,619 with a range of 1,086 to 822,149.

Species’ richness was estimated from each sample using the rarefied number of species and species’ diversity was estimated by Simpson’s diversity index using the vegan package in R (Oksanen et al., 2015). Species’ richness and diversity were tested for association with sample substrate (bark, soil, must) using an ANOVA and pairwise differences were assessed using Tukey’s method. For the fermentation time-course, species diversity and richness did not change linearly over time and so we fit a linear model to the ranked order of richness and diversity from each fermentation. Nonmetric multidimensional scaling was implemented using the metaMDS function in the vegan package of R with 20 starting points based on Bray-Curtis dissimilarity among samples.

### Wine phenotypes

Strains were grown in 200 µl of complete medium (2% glucose, 2% yeast nitrogen base with ammonium sulfate) overnight in 96-well plates. Strains were then resuspended 1:20 in complete medium with sulfite, copper, ethanol, tartaric acid or unaltered and grown for 48 hrs without shaking at 30°C in 96-well plates. Sulfite medium was 0.7 and 1.5 mM Na2SO3 buffered to a pH of 3.5 with tartaric acid. Low pH was 5 mg/mL of tartaric acid, copper medium was 0.5 mM and 1.0 mM copper sulfate, ethanol medium was 6% and 10% (v/v) ethanol. These concentrations were selected based on preliminary assays to distinguish North American oak and commercial wine strains. Cell density (OD600) was measured (iEMS plate reader, Thermo Lab Systems, Helsinki, Finland) at 0, 19, 24, 36 and 48 hrs after treatment. Data for certain time-points, 1.4% of all the data, was interpolated due to plate reader malfunction: two of the plates for sulfite treatment at 19 hrs, one plate for tartaric acid at 24 hrs, and one plate at 48 hrs for the no stress control. Data were interpolated by taking the average of the prior and subsequent time-points. The phenotype of each strain was measured by the area under the growth curve (AUC) and we used the average AUC when growth was measured under two different stress concentrations. Commercial strains were obtained from yeast distributors. North American oak tree strains were those from Hyma et al. (2013) from which we excluded *S. cerevisiae* and *S. paradoxus* strains closely related to European strains.

### Reanalysis of North American samples

From the raw data of Hyma et al. (2013) we analyzed 187 *S. cerevisiae* and 240 *S. paradoxus* isolates from 977 oak-and 492 grape-associated samples for which the same enrichment medium was used. The frequency of each species was fit to a logistic model with terms for state (MO, OR), location type (vineyard, non-vineyard), source (oak, grapes) and year of isolation (2008, 2009). Significant terms were identified by dropping single terms and comparing models using a likelihood ratio test.

## Results

### Isolation of Saccharomyces yeasts

To characterize the distribution and abundance of *Saccharomyces* species we sampled 1,233 substrates from 7 vineyards and 4 non-vineyard locations in Slovenia between July of 2013 and April of 2014. Substrates were primarily from oak trees (66%) and grapevines (24%). The remaining samples were from wine cellars, must, fruit, insects and other plant material (Table S3). Following enrichment of the samples, we isolated 869 strains and distinguished *Saccharomyces* species from one another and from non-*Saccharomyces* species (Methods). Our sample yield was highest for non-*Saccharomyces* species (28%), followed by *S. paradoxus* (23%), *S. cerevisiae* (12%) and *S. kudriavzevii* (2.1%) (Table S3).

To test whether enrichment at higher temperature increased our recovery of *S. cerevisiae*, we split 518 of the samples into enrichments at room temperature and 37ºC. High temperature enrichments yielded a higher ratio of *S. cerevisiae* relative to *S. paradoxus* strains (29:1 compared to 81:123, Fisher’s Exact Test P < 0.01). However, substantially fewer high temperature enrichments yielded yeast (11%) compared to those at room temperature (80%) due to proliferation of bacteria (Fisher’s Exact Test P < 0.01). The higher ratio of *S. cerevisiae* to *S. paradoxus* strains from high temperature enrichments was not a primary consequence of temperature since most of the corresponding low temperature enrichments from the same sample yielded the same species and in only four cases was *S. paradoxus* isolated at the low temperature when *S. cerevisiae* was isolated at the high temperature. To avoid potentially redundant samples, we removed the 55 high temperature isolates from the remainder of the analysis.

### Species abundance differs by source, geographic location and time of year

As a proxy for species abundance, we compared rates of isolation from 1,055 samples associated with oak trees within the vineyard (467), oak trees outside the vineyard (316), and grapevines (272). While we found no differences between vineyard and non-vineyard locations, *S. cerevisiae* and non-*Saccharomyces* yeasts were more prevalent on grapevine-compared to oak-associated substrates and *S. paradoxus* was depleted (Figure 1, Table S4). We also found variation across sampling locations for both *S. cerevisiae* and *S. paradoxus*, but not for non-*Saccharomyces* yeast as a group (Table S4 and S5). One apparent outlier was an abandoned castle on a hill overlooking the town of Vipava; it was the only non-vineyard location with more *S. cerevisiae* than *S. paradoxus* isolates. However, removing this location still yielded equivalent ratios of *S. cerevisiae* to *S. paradoxus* from vineyard oaks and non-vineyard oaks (odds ratio [95% confidence interval] = 0.23 [0.16,0.32] and 0.13 [0.052,0.26], respectively, P = 0.172).

**Figure 1.**
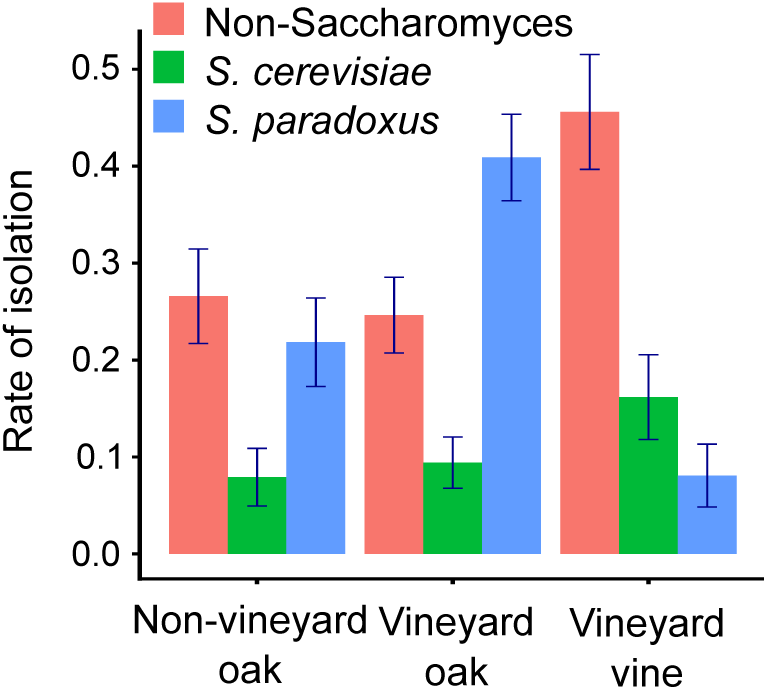
Rates of isolation depend on sample source. The sampling frequency of each species is shown for oak-associated samples within and outside of vineyards, and for grapevine-associated samples.

Time of year influenced the sampling rate of all species except for *S. cerevisiae* (Table S3). In September during harvest time we found the lowest rates of *S. paradoxus* and the highest rates of non-*Saccharomyces* yeast (Table S4). Except for one isolate from April, all *S. kudriavzevii* strains were obtained in October, mostly from non-vineyard oak samples.

### *Saccharomyces* abundance within the oak-and grape-associated microbiomes

To quantify the relative abundance of *Saccharomyces* and other yeast species without enrichment, we performed ITS1 sequencing on 20 vineyard samples of oak bark and soil and 20 samples from uninoculated wine must. Bark and soil samples contained more species than must samples (Tukey P < 0.001), but there was no difference in Simpson’s diversity index, which measures the skew towards one or a small number of abundant species (Tukey P > 0.05, Table S6). However, species’ relative abundance differed across sample substrates. At a broad taxonomic level, five out of six classes with overall abundance above 5% differed in frequency among must, bark and soil samples (Figure 2, ANOVA P < 0.01), with the one exception being *Dothideomycetes* which were abundant in all samples. Must samples were enriched for *Saccharomycetes* and *Leotiomycetes*, with the most common species being *S. cerevisiae* and the grape pathogen *Botryotinia fuckeliana*, respectively. Bark samples were enriched for *Lecanoromycetes*, with the most common species being the lichen *Physciella chlorantha*, and soil samples were enriched for *Agaricomycetes*, with the most common being the mushroom *Russula fragilis*. Multidimensional scaling of species’ abundance also distinguished must from bark and soil samples, the latter two of which were more similar to one another (Figure S1).

**Figure 2.**
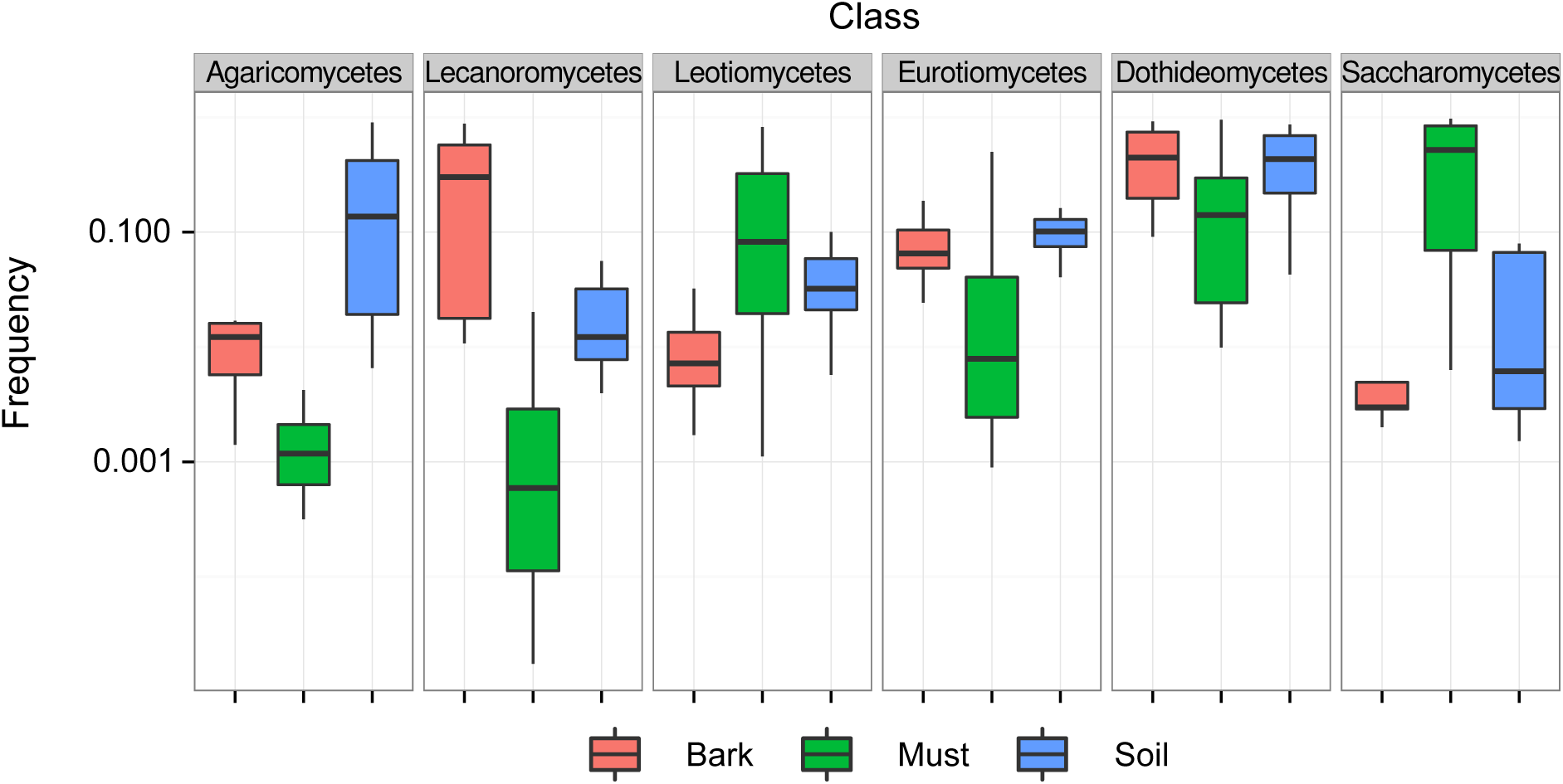
Frequency of abundant taxonomic classes differs across samples. Boxplots are shown for six abundant classes (> 5%) grouped by substrate from which they were obtained.

The frequency of *Saccharomyces* species was highly variable across samples (Figure 3). Both *S. cerevisiae* and *S. paradoxus* were rare in soil and bark samples, averaging 6.9×10^−4^ for *S. cerevisiae* and 6.6×10^−5^ for *S. paradoxus*. The two species were more variable in must samples, with *S. paradoxus* constituting up to 82% and *S. cerevisiae* up to 87% of identified species (Table S6). Although *S. cerevisiae* tended to have a higher frequency than *S. paradoxus* across all samples (Mann-Whitney test, P = 6.7×10^−5^), there was no significant difference among substrates in the relative abundance of *S. cerevisiae* to *S. paradoxus* (Kruskal Wallis test, P = 0.085). Another *Saccharomyces* species found, *S. kudriavzevii*, was only present in a single sample (SM56) at a frequency of 4.8×10^−5^.

**Figure 3.**
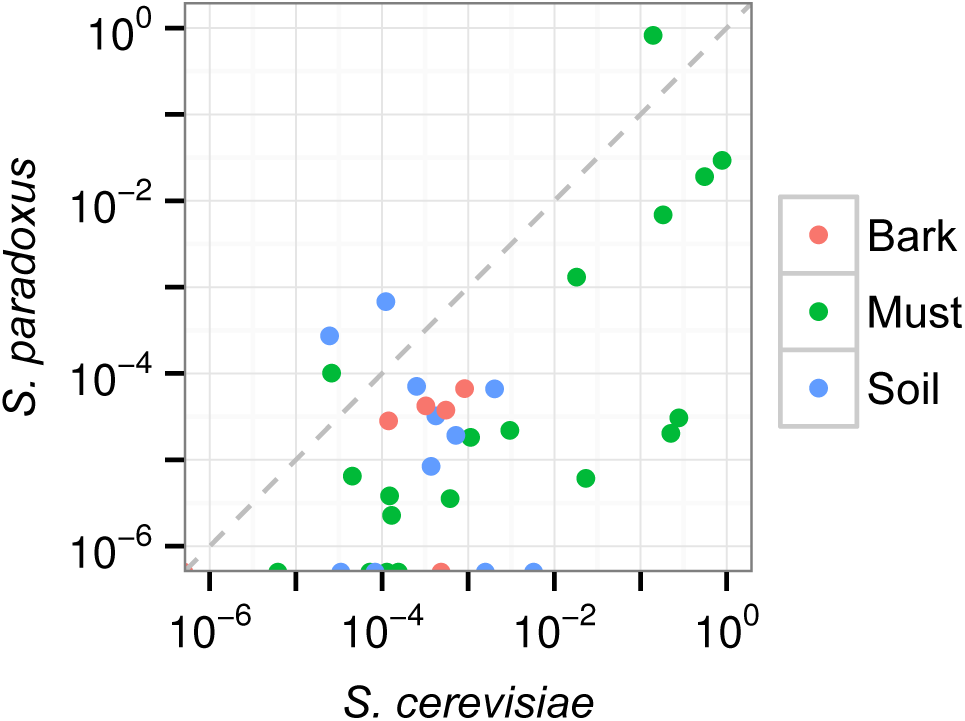
Frequency of *S. cerevisiae* and *S. paradoxus* in soil, bark and must samples. Points shown on the x-axis had no *S. paradoxus* counts.

### *S. cerevisiae* increases in abundance during wine fermentations

To capture changes in temporal dynamics that occur during fermentation we obtained juice from the Carga estate and grape pomace from the Burja estate (previously a part of Sutor), and carried out triplicate fermentations in the lab, taking 8–10 samples over 14–17 days of fermentation. In parallel, we sampled the same pomace from the Burja estate that was being fermented in the Burja winery cellar.

Species’ richness and Simpson’s diversity index decreased over time for both the Burja (P = 0.02 and P = 3.3×10^−5^, respectively) and Carga (2.9×10^−6^ and 4.1×10^−9^, respectively) experimental fermentations but not for the Burja cellar fermentation (P > 0.05, see Methods). Initial richness and diversity of the experimental fermentations was within the range of the 20 must samples (Table S7) and primarily consisted of *Saccharomycetes* (Figure 4). By the end of fermentation, only three species were above 5%: *S. cerevisiae*, *Starmerella bacillaris* and *Botryotinia fuckeliana*. Interestingly, *S. paradoxus* was absent from the Carga fermentation and was only present at an initial frequency of 3.8×10^−4^ in the Burja fermentation. In comparison, *S. cerevisiae* was at an initial frequency of 8.7×10^−3^ and 8.2×10^−2^ in the Carga and Burja fermentations, respectively.

**Figure 4.**
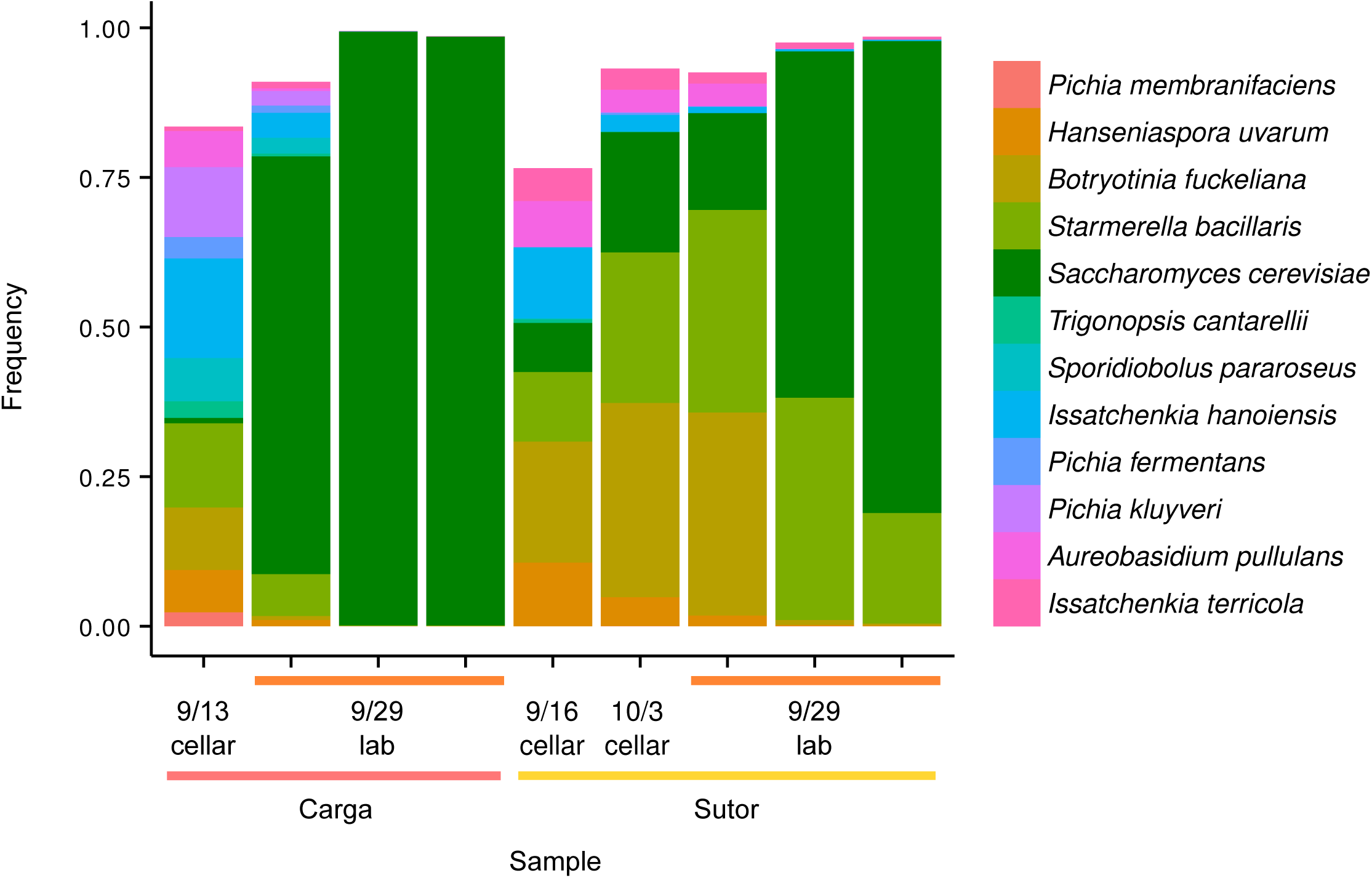
Changes in species abundance during fermentation. Twelve species with at least 5% abundance at one time-point are shown for Burja and Carga fermentations at the start (cellar) and end (lab, 3 replicates) of each time-course. Also shown is a Burja cellar sample at the start and end of fermentation. Counts of *Saccharomyces boulardii* were included in those of *S. cerevisiae*.

### European strains of *S. cerevisiae* have acquired resistance to stresses applied during wine making

The presence of both *S. cerevisiae* and *S. paradoxus* within vineyard and wine must suggests *S. paradoxus* should often make it into wine fermentations. However, the Carga and Burja fermentations along with previous work (Bokulich et al., 2014b; David et al., 2014; Gayevskiy and Goddard, 2012; Pinto et al., 2014, 2015; Setati et al., 2012; Taylor et al., 2014; Wang et al., 2015) indicate that it may not often be a major contributor to fermenting wine must. One potential cause for a shift in the relative abundance of these two species going from the vineyard into the winery is the addition of copper and sulfites to the grape must. Indeed, sulfites were added to both the Carga and Burja musts before being brought to the lab. Previous studies have shown wine strains are particularly resistant to copper and sulfites (Liti et al., 2009; Warringer et al., 2011). To characterize sensitivity to the wine making environment among our isolates we measured the growth profiles of 168 *S. cerevisiae* and 263 *S. paradoxus* from Slovenia in comparison to a set of 35 reference commercial wine strains, 29 North American oak tree strains, and 34 North American *S. paradoxus* strains (Hyma and Fay, 2013). As a control we measured growth in the absence of stress and in the presence of ethanol, which has not been reported to differ between the two species.

As expected, North American *S. cerevisiae* strains are more sensitive than commercial wine strains to sulfites, copper and low pH, but not high ethanol (FDR < 0.01, Figure 5, Figure S2, Table S8). Slovenian *S. cerevisiae* strains are resistant to sulfites, copper and low pH; more so than North American *S. cerevisiae* or *S. paradoxus* (FDR < 0.01 Table S8). This high level of resistance of Slovenian *S. cerevisiae* strains is indistinguishable from that of commercial wine strains (FDR > 0.01).

**Figure 5.**
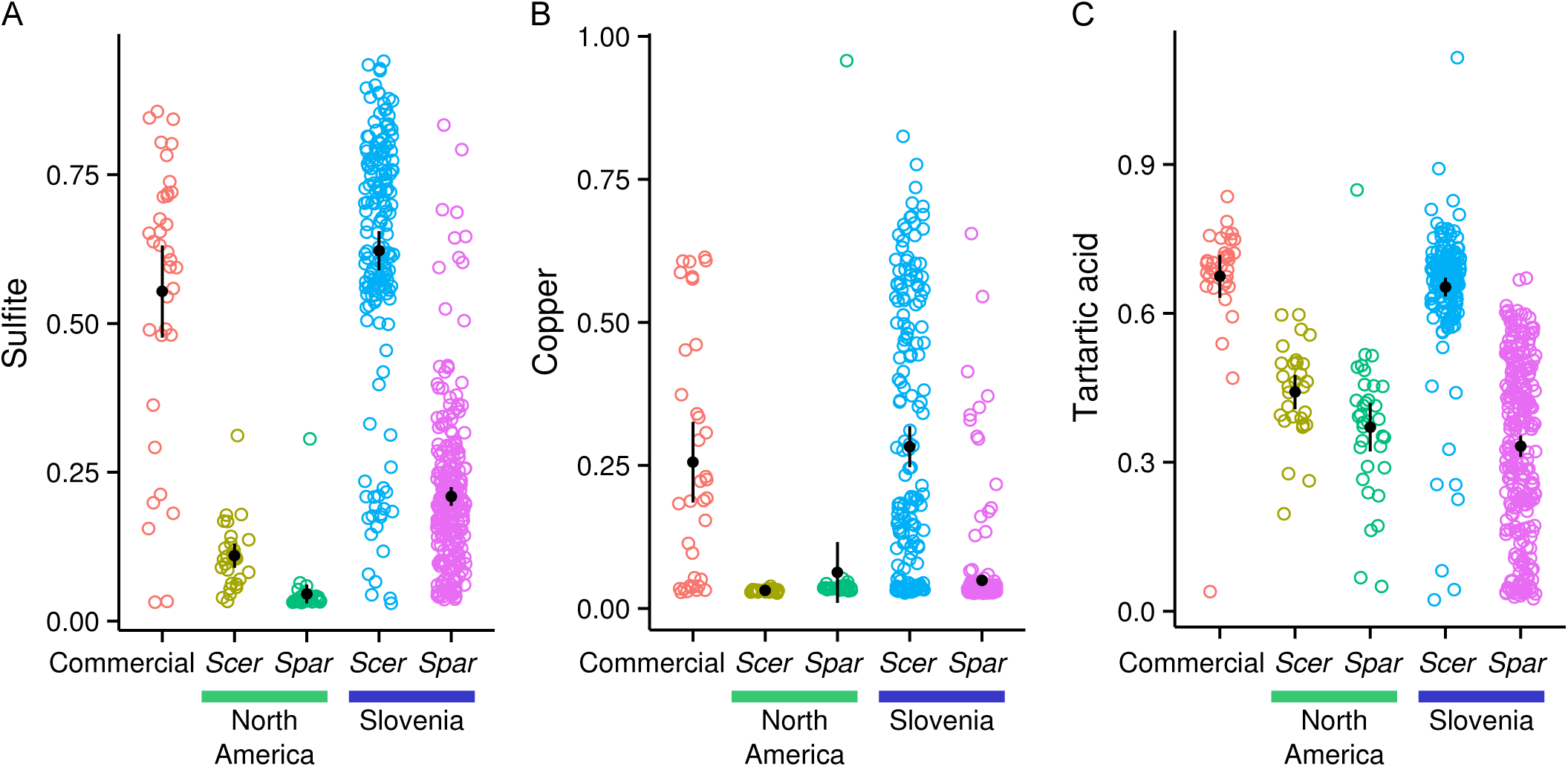
Slovenian *S. cerevisiae* strains are resistant to sulfite, copper and tartaric acid. Growth rates (area under the growth curve) in the presence of sulfite (A), copper (B) and tartaric acid (C) relative to the absence of stress for *S. cerevisiae* (Scer), *S. paradoxus* (Spar) and commercial wine strains. Black circles and bars represent the mean and its 95% confidence interval.

### Discussion

Strains of *S. cerevisiae* have been widely used for the production of beer, bread, wine and other human-associated fermentations (Sicard and Legras, 2011). Its sibling species, *S. paradoxus*, is rarely associated with human fermentations (Boynton and Greig, 2014) but is a strong fermenter and is competitive with *S. cerevisiae* in grape juice (Williams et al., 2015). The distinction between these two species is particularly well defined in Europe, where *S. cerevisiae* is most often isolated from vineyards whereas *S. paradoxus* is most often isolated outside of vineyards.

In this study we used intensive sampling and microbial profiling to show that there is not a clear cut difference in the abundance and distribution of these two species within Slovenian vineyards and forests. Similar to North American vineyards and forests (Hyma and Fay, 2013), the two species can occur sympatrically in Europe and we find they only differ in their relative abundance: *S. paradoxus* is more abundant on oak tree-associated substrates and *S. cerevisiae* is more abundant on grapevine-associated substrates. Although there are likely many factors, discussed below, that contribute to variation in the relative abundance of these two species, our results support the idea that current wine making practices greatly enrich *S. cerevisiae* within the vineyard via the acquisition of multiple traits by European wine strains.

### Is *S. paradoxus* rare within vineyard environments?

Our results based on both enrichment and microbiome analysis indicate that *S. paradoxus* is not excluded from vineyard environments, including wine must, and can be as abundant as *S. cerevisiae*. While our findings differ from those of prior studies, multiple factors influence the relative abundance of these two species and may explain these differences.

### Distinguishing *Saccharomyces* species

Because the *Saccharomyces* species were not clearly delineated until the 1990s (Naumov, 1996; Naumov et al., 1992), early work on yeasts present within vineyard and wine fermentation may not have distinguished between *S. cerevisiae* and *S. paradoxus*. Even so, *S. cerevisiae* remains the predominant yeast isolated from European vineyards, e.g. in Spain (Cordero-Bueso et al., 2011), Portugal (Schuller et al., 2005), Italy (Di Maio et al., 2012; Stefanini et al., 2012) and France (Valero et al., 2007). With the exception of Mediterranean regions where *S. cerevisiae* is found to co-occur with *S. paradoxus* on trees (Almeida et al., 2015; Sampaio and Gonçalves, 2008), *S. paradoxus* is the predominant *Saccharomyces* species isolated from forest environments (Bozdag and Greig, 2014; Glushakova et al., 2007; Johnson et al., 2004; Koufopanou et al., 2006; Kowallik et al., 2015), reinforcing the notion that *S. paradoxus* is a wild yeast and absent from vineyards (Boynton and Greig, 2014).

More recently, the diversity of yeasts present within fermenting wine have been examined by direct sequencing of the wine microbiome (Bokulich et al., 2014b; David et al., 2014; Gayevskiy and Goddard, 2012; Pinto et al., 2014, 2015; Setati et al., 2012; Taylor et al., 2014; Wang et al., 2015). While these microbiome studies have not reported *S. paradoxus* within the wine must, certain methods of analysis do not distinguish it from *S. cerevisiae*. One common practice is the representation of closely related sequences at the level of 97–99 percent identity by operational taxonomic unites (OTUs). In the UNITE database (Kõljalg et al., 2013), these groups are termed species hypothesis (SH) and do not distinguish *S. cerevisiae* from *S. paradoxus* even though they are readily distinguishable by their ITS1 sequence (McCullough et al., 1998). Thus, the absence of reports of *S. paradoxus* within vineyards and wine must may be partly attributed to not specifically distinguishing it from *S. cerevisiae*.

### Variation across ecological niches

In our samples we found the relative abundance of *S. cerevisiae* to *S. paradoxus* is related to habitat, grapevines or oak trees, but not whether the oak trees occur within or outside of vineyards. As such, it is not surprising that we also isolated *S. paradoxus* from grapevine-associated substrates and that the habit surrounding vineyards is relevant to the microbial community colonizing grapevines and being incorporated into wine must (Bokulich et al., 2014b; Knight et al., 2015; Setati et al., 2012).

In contrast to our enrichment samples, our microbial profiling experiments detected no significant differences in the relative abundance of *S. cerevisiae* to *S. paradoxus* among must, bark or soil samples; *S. cerevisiae* was found to be uniformly more abundant. This difference could be a consequence of the low frequency of *S. cerevisiae* and *S. paradoxus* in most bark and soil samples, close to the detection limit of 10^−4^ to 10^−5^ determined by the number of sequence reads per sample. Furthermore, the sample size was small relative to the number of enrichment samples and we observed substantial sample to sample variation in the relative abundance of the two species. Finally, we cannot exclude the possibility that our enrichment process generated a biased representation of species abundance. The presence of other microbes in a sample can influence yeast growth (Kowallik et al., 2015) and so it is possible that differences in sample abundance occurred because *S. cerevisiae* and *S. paradoxus* differ in their ability to compete with microbes that are not evenly distributed across oak and grapevine habitats.

### Local geographic and temporal variation

By design we sampled multiple vineyards to help ensure our results were reflective of Slovenian vineyards and the Vipava valley. By necessity we sampled multiple times during the year, with the majority of samples being collected before (July) and after (October) harvest. Both location and time of year are associated with the relative abundance of *S. cerevisiae* to *S. paradoxus*. The most interesting deviation within our sampling locations was an abandoned castle on a hill in Vipava. While the castle itself is old (13^th^ Century) the oak forest surrounding it consist of young oak trees, approximately 10–20 cM in diameter at the base of the tree. While not optimal for characterizing species abundance in European forests, this location highlights the importance of fine-scale variation and historical context in sampling locations.

While our finding of sympatric *S. cerevisiae* and *S. paradoxus* within and outside of vineyards in Slovenia may be a regional finding, it is consistent with certain studies. *S. paradoxus* was found in a vineyard in a region of Croatia approximately 150 km East of Vipava (Redzepović et al., 2002), and wild populations of *S. cerevisiae* associated with Mediterranean oaks have been isolate from Southern Europe (Almeida et al., 2015).

### Global geographic variation

Our work establishes *S. paradoxus* as part of the vineyard environment, at least in Slovenia. While this raises the possibility that it may also occur in vineyards outside of the Balkans, *S. paradoxus* has thus far only been isolated from North American vineyards (Hyma and Fay, 2013). Because Hyma and Fay (2013) did not report rates of isolation, we analyzed the raw data for comparison with our results from Slovenia.

Similar to Slovenia, numerous isolates (130) of *S. paradoxus* were obtained from vineyards (Table S9). Highlighting the importance of geographic variation, *S. paradoxus* was almost exclusively isolated from both vineyard and forest locations in Oregon. Yet even accounting for geographic variation, *S. paradoxus* was less abundant in vineyards (OR = 0.55, P = 0.03) and not significantly different from *S. cerevisiae* when comparing oak versus grapevine samples (OR = 0.83, P = 0.73, Table S10). Thus, while both the Slovenian and United States samples show evidence of geographic variation, the United States samples differ by sample location (vineyard vs. forest) rather than sample source (grapevine versus oak). However, it should be noted that in the US only 22 isolates were obtained from grapes and 16 of these were *S. cerevisiae*.

### Transition into the winery and competition during fermentation

Similar to other studies of microbial diversity (Combina et al., 2005; Mercado et al., 2007; Pinto et al., 2015), we found a diverse fungal community from harvested and/or pressed grapes before or at the initial stages of fermentation followed by a rapid decline in diversity as *S. cerevisiae* became the dominant species. While the grape must community was distinct from oak bark and soil communities, the community was also quite variable at the species level. This variability could be related to any number of differences in location, method and time of harvest or contact with winery equipment. Along with overall variation in the grape must microbiome, the relative abundance of *S. cerevisiae* and *S. paradoxus* also varied, with one sample of must from pressed grapes containing 82% *S. paradoxus* and only 14% *S. cerevisiae*.

One limitation of our grape/must samples is that we did not control for sulfite or other treatments of the grapes or must before sampling. Although we sampled from wineries that carry out autochthonous fermentations, the vineyards also spray copper sulfate as a fungicide and use sulfites to inhibit the growth of bacteria and other microorganisms. Indeed, sulfites were added by the wineries prior to deriving wine fermentations in the lab. Such treatments very likely alter initial microbial diversity or their dynamics during fermentation to wine (Bokulich et al., 2014a). Our observation that unlike *S. cerevisiae*, *S. paradoxus* does not become common during wine fermentation can be explained by differences in resistance to copper, sulfite or low pH. However, these aspects of the experimental wine fermentation were not controlled and so further work will be needed to distinguish them from other factors that could be involved.

Even after being brought into the lab we observed substantial variation in species abundance during fermentation to wine. The most notable difference was the maintenance of much higher levels of diversity in the Burja wine fermented at large volume in the cellar and in one of our laboratory replicates as compared to the other two replicates carried out in the lab. The cellar fermentation could be different due to a slower rate of fermentation, larger volume or lower temperature, but could also be due to our mixing the lab fermentation prior to every sample taken.

### Resistance to copper, sulfites and tartaric acid distinguishes Slovenian strains of *S. cerevisiae* and *S. paradoxus*

Prior work has shown that both resistance to copper and sulfites are common in wine strains compared to oak strains of *S. cerevisiae* and *S. paradoxus* (Fay et al., 2004; Liti et al., 2009; Pérez-Ortín et al., 2002; Strope et al., 2015; Warringer et al., 2011; Yuasa et al., 2004), as might be expected given their frequent use in vineyards and wineries. Our phenotypic analysis of Slovenian yeast adds resistance to low pH to these two previously characterized “domestication” phenotypes and shows that these phenotypes differentiate vineyard isolates of *S. paradoxus* from European but not North American *S. cerevisiae*. Thus, we can conclude that the sensitivity of *S. paradoxus* to copper, sulfite and low pH is not because *S. paradoxus* is absent from vineyards and hasn’t had the opportunity of facing selective pressures that are common in the vineyard environment.

The acquisition of copper and sulfite resistance in wine strains has been extensively studied and is known to be primarily caused by changes at *CUP1* (Adamo et al., 2012; Chang et al., 2013; Fogel and Welch, 1982; Strope et al., 2015; Zhao et al., 2014) and *SSU1* (Goto-Yamamoto et al., 1998, 1; Nardi et al., 2010; Yuasa et al., 2004, 1; Zimmer et al., 2014), respectively. However, the relationship between resistance to sulfite and tartaric acid is less clear. Sensitivity to sulfite was measured at a pH of 3.5 since there is little of the active agent sulfur dioxide at higher pH (Casalone et al., 1992). Sensitivity to low pH was measured by adding tartaric acid, since it is abundant in grapes (Kliewer et al., 1967). While resistance between the two is correlated (r^2^ = 0.44), resistance to tartaric acid only explains 3% of variation in sulfite resistance once differences among major groups (64% of variation, Figure 5) are accounted for.

### Conclusions

The history and origins of wine strains has begun to emerge with detailed studies of *S. cerevisiae* in comparison to its closest known relative *S. paradoxus* (Boynton and Greig, 2014). While certain aspects of these two species are notably different, they are sympatric in North American forests (Hyma and Fay, 2013; Sniegowski et al., 2002) and our present results demonstrate that they can inhabit the same vineyard environments. Thus, *S. paradoxus* may be similar to *S. cerevisiae* in its opportunistic colonization of certain environments (Goddard and Greig, 2015). However, one of the fundamental differences between these two species is the higher diversity and stronger geographic structure of *S. paradoxus* compared to *S. cerevisiae* (Liti et al., 2009). Not only is the spread of European wine strains relevant to *S. cerevisiae* population structure (Fay and Benavides, 2005), but there is now also evidence for the spread of wild oak populations of *S. cerevisiae* based on the clonal relatedness of isolates from North America and Japan (Almeida et al., 2015; Hyma and Fay, 2013). Thus, the current sympatric relationship between *S. cerevisiae* and *S. paradoxus* in Slovenian vineyards, and perhaps North American forests, may be a relatively recent development. Further elucidation of the history and relationship between these two species will have to meet the challenge of geographic and temporal heterogeneity while accounting for the historic use or vegetation of the habitats sampled. With sufficient fortitude or luck we may be able to better define the vectors and environmental reservoirs, humans-associated or otherwise, pertinent to these closely related but differentially exploited species.

## Author Contributions

Conceived and designed the experiments: SD, LB, JP and JF. Collected and analyzed data: SD, PL, HV, LB, JP and JF. Wrote the paper: SD and JF.

## Acknowledgements

This work was supported by RS-MIZS and European Regional Development Fund Research (the Creative Core programme “AHA-MOMENT”, contract no. 3330-13-500031), by the Slovenian Research Agency (ARRS J4-4300 and BI-US/13-14-028) and the National Institutes of Health (GM080669). We are particularly grateful to the owners and staff of the Carga, Guerilla, Sveti Martin, Sutor, Burja, University of Nova Gorica, Rencelj and Tilia estates and wineries for allowing us to sample their vineyards, wine cellars and wine.

## Supplementary Material

**Figure S1.**
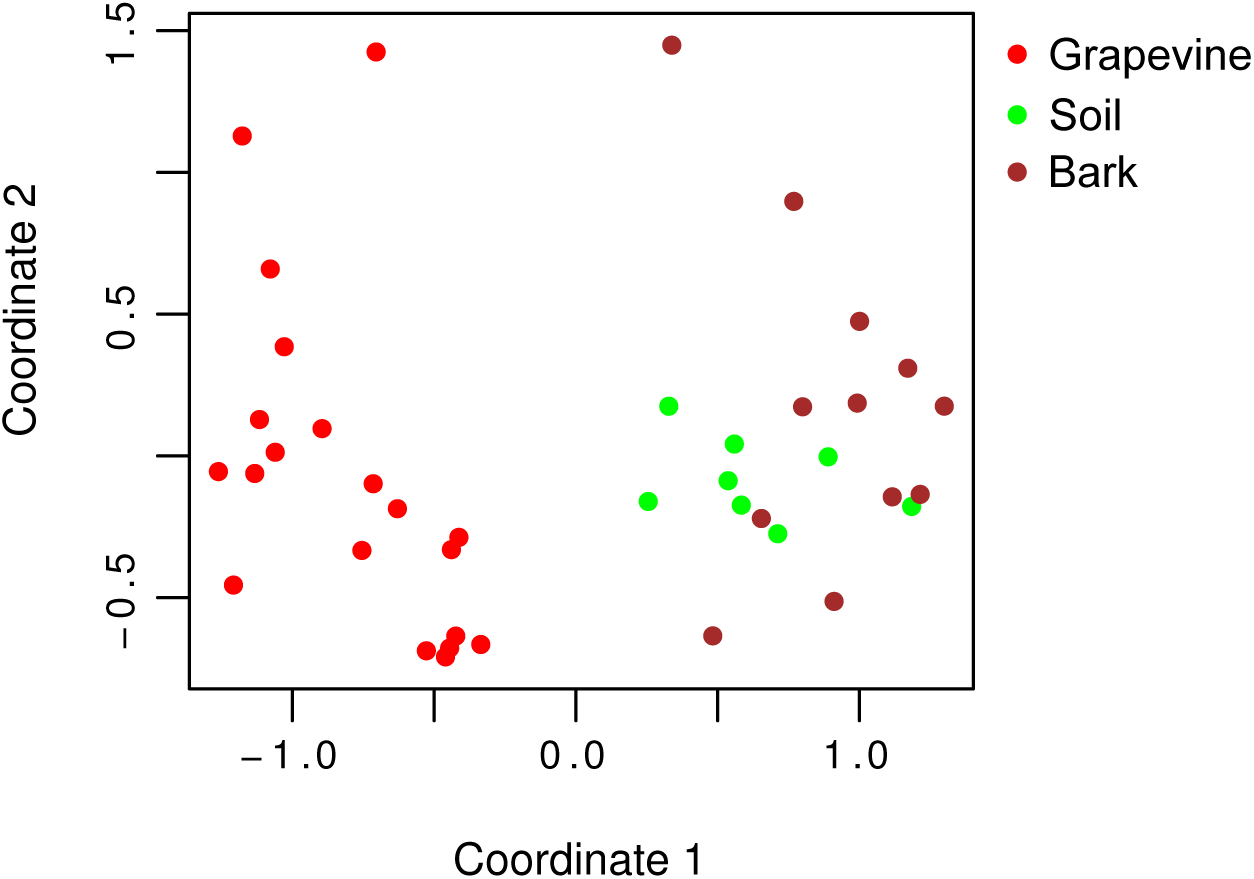
Multidimensional scaling of microbiome samples. First and second coordinates are from non-metric multidimensional scaling using Bray-Curtis dissimilarity.

**Figure S2.**
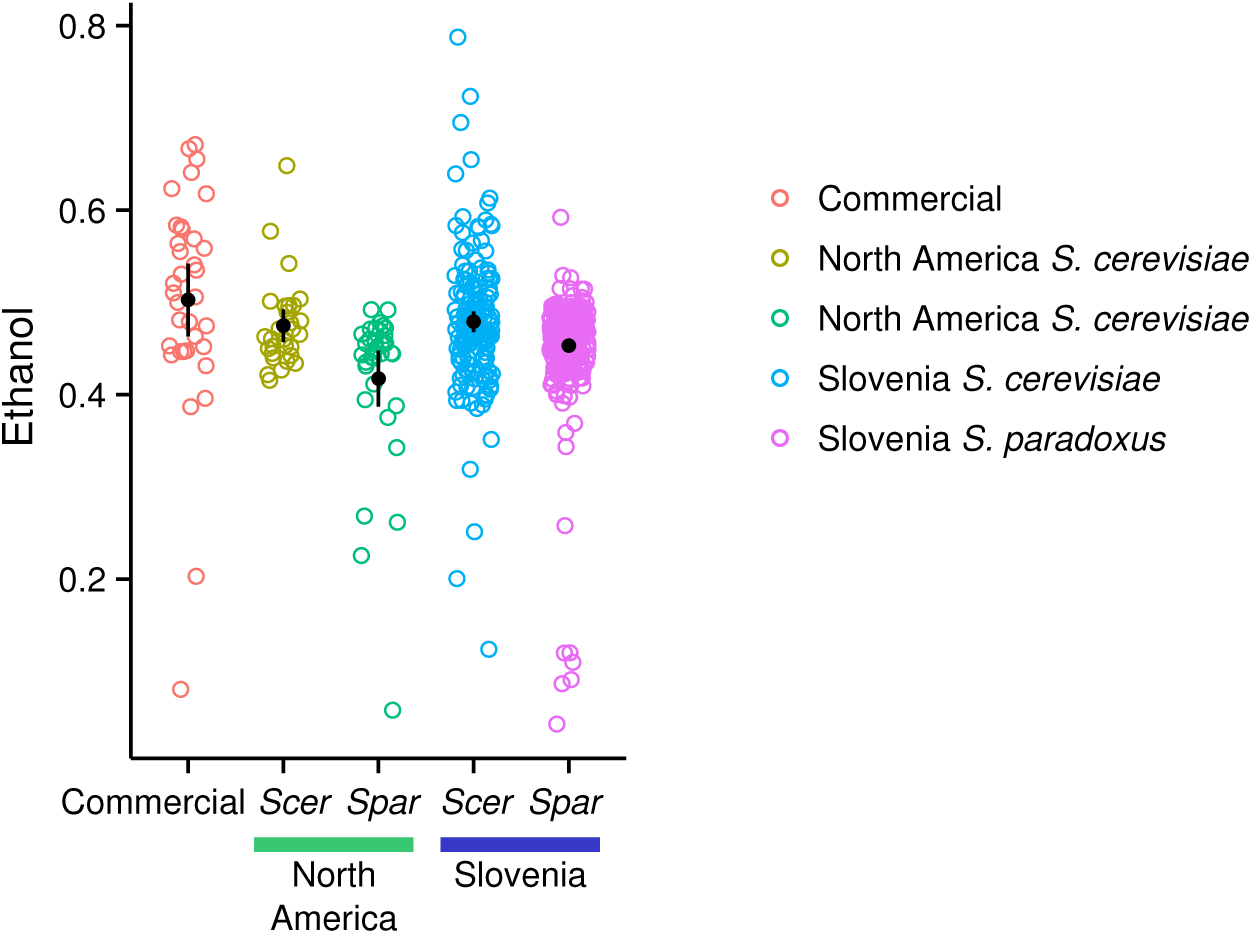
Slovenian and North American strains’ resistance to ethanol. Growth rates (AUC) in the presence of ethanol relative to its absence. Black circles and bars represent the mean and its 95% confidence interval.

Data File S1. Samples used for enrichment and species identified.

Data File S2. Species counts across microbiome samples.

Data File S3. Phenotype data.

Data File S4. Samples analyzed from North America.

Data File S5. Microbiome raw reads and metadata (this data file will be submitted to dryad once the manuscript has been accepted in accordance with dryad policy).

Data File S6. Supporting tables S1-10.

**Table S1.** Sampling locations.

| Name | Type | Location | Latitude | Longitude | Samples |
| --- | --- | --- | --- | --- | --- |
| Castle | Forest | Vipava | 45.850334 | 13.964775 | 110 |
| Lijak | Forest | Lijak | 45.959051 | 13.719193 | 82 |
| Nanos | Forest | Nanos | 45.796672 | 14.002717 | 68 |
| Sinji Vrh | Forest | Sinji Vrh | 45.907952 | 13.935454 | 97 |
| Carga | Vineyard | Brda | 46.02785 | 13.518732 | 200 |
| Guerila | Vineyard | Planina | 45.84935 | 13.909625 | 101 |
| Rencelj | Vineyard | Dutovlje | 45.752125 | 13.830302 | 25 |
| Sutor | Vineyard | Podraga | 45.811386 | 13.943568 | 160 |
| Sveti Martin | Vineyard | Brje | 45.860662 | 13.812358 | 114 |
| Tilia | Vineyard | Dobravlje | 45.892136 | 13.833961 | 120 |
| UNG | Vineyard | Manče | 45.816556 | 13.941429 | 156 |

**Table S2.**
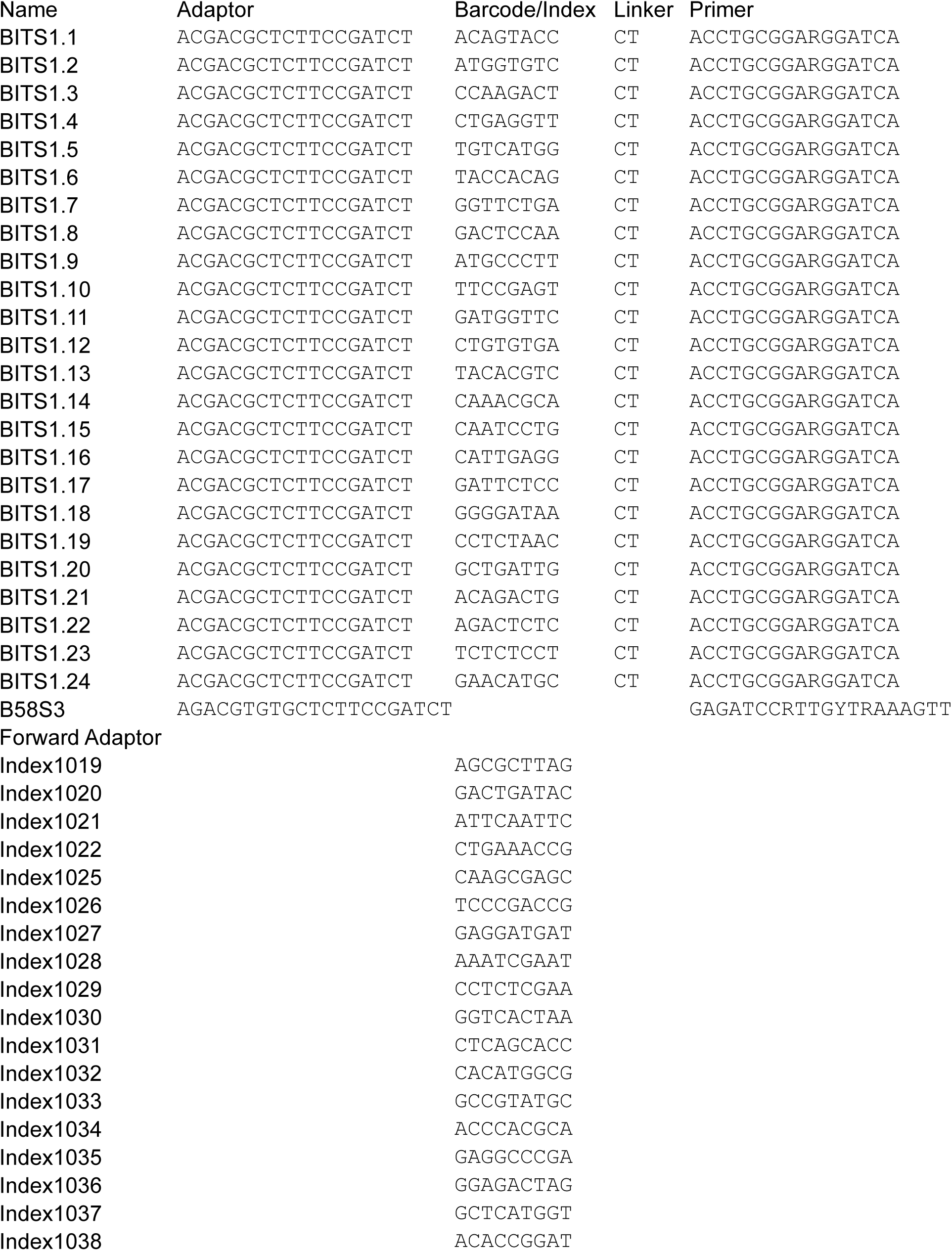
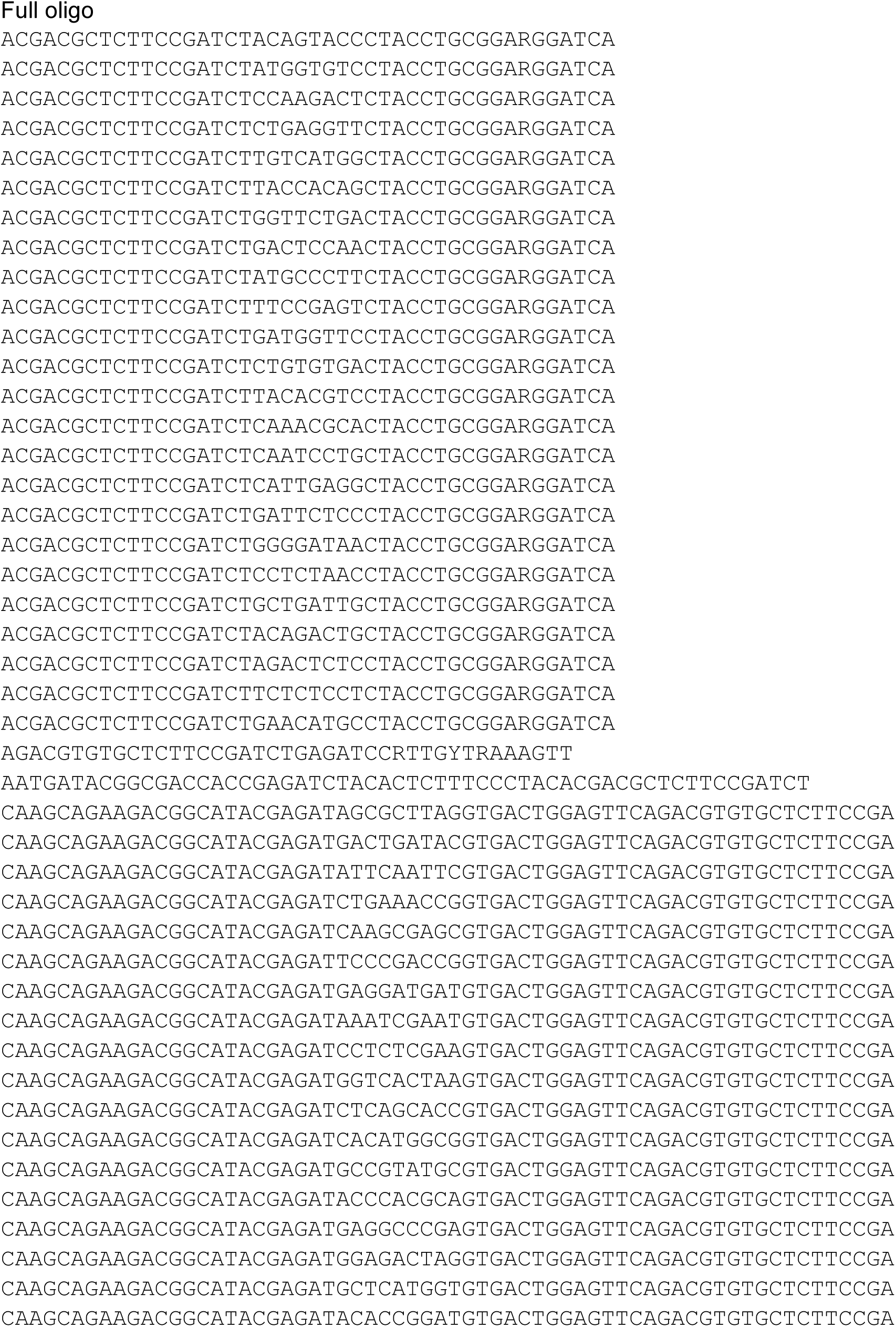
Primers used for microbiome analysis.

**Table S3.** Isolated species by sample source.

| Source | Samples | *S. cerevisiae* | *S. paradoxus* | *S. kudriavzevii* | Non-Saccharomyces | Failed identification |
| --- | --- | --- | --- | --- | --- | --- |
| Cellar | 19 | 18 | 0 | 0 | 1 | 0 |
| Fruit | 16 | 3 | 2 | 0 | 5 | 0 |
| Insect | 47 | 1 | 1 | 0 | 11 | 6 |
| Must | 36 | 14 | 0 | 0 | 7 | 2 |
| Oak | 809 | 69 | 260 | 25 | 199 | 26 |
| Vine | 294 | 46 | 22 | 1 | 125 | 19 |
| Yeast | 2 | 2 | 0 | 0 | 0 | 0 |
| Other | 10 | 0 | 2 | 0 | 2 | 0 |
| All | 1233 | 153 (12.3%) | 287 (23%) | 26 (2.1%) | 350 (28%) | 53 (4.2%) |

Other = other types of plant material, Yeast = commercial yeast used by vineyard

**Table S4.**
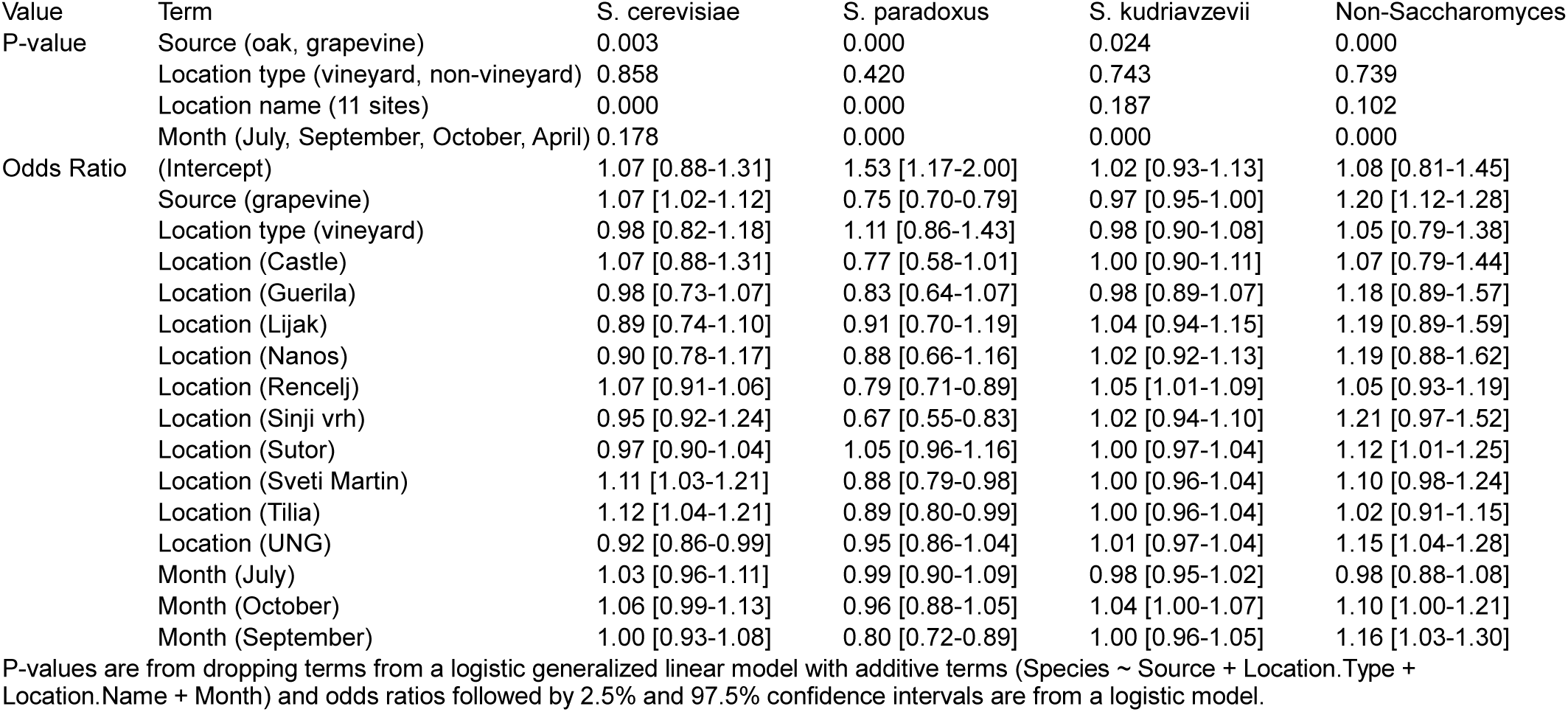
Factors influencing rates of isolation.

**Table S5.**
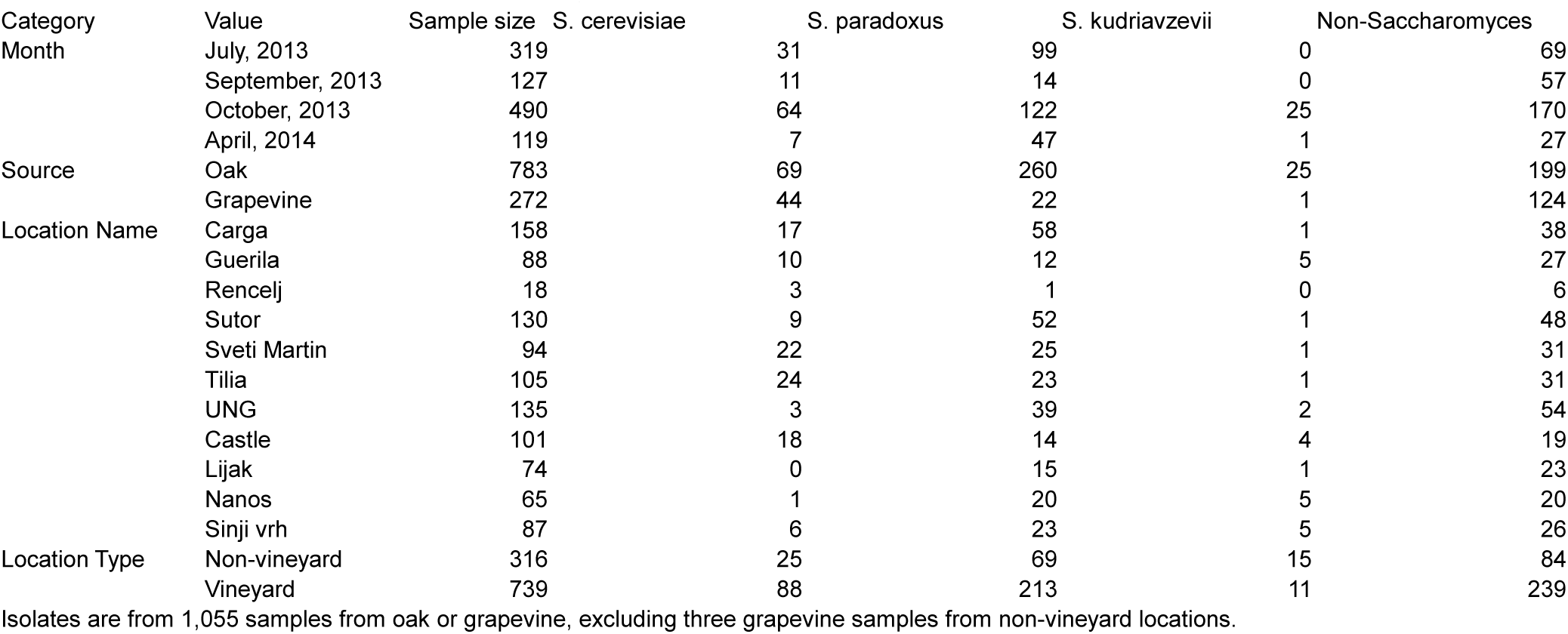
Sample size and number of isolates by month, source and location.

**Table S6.**
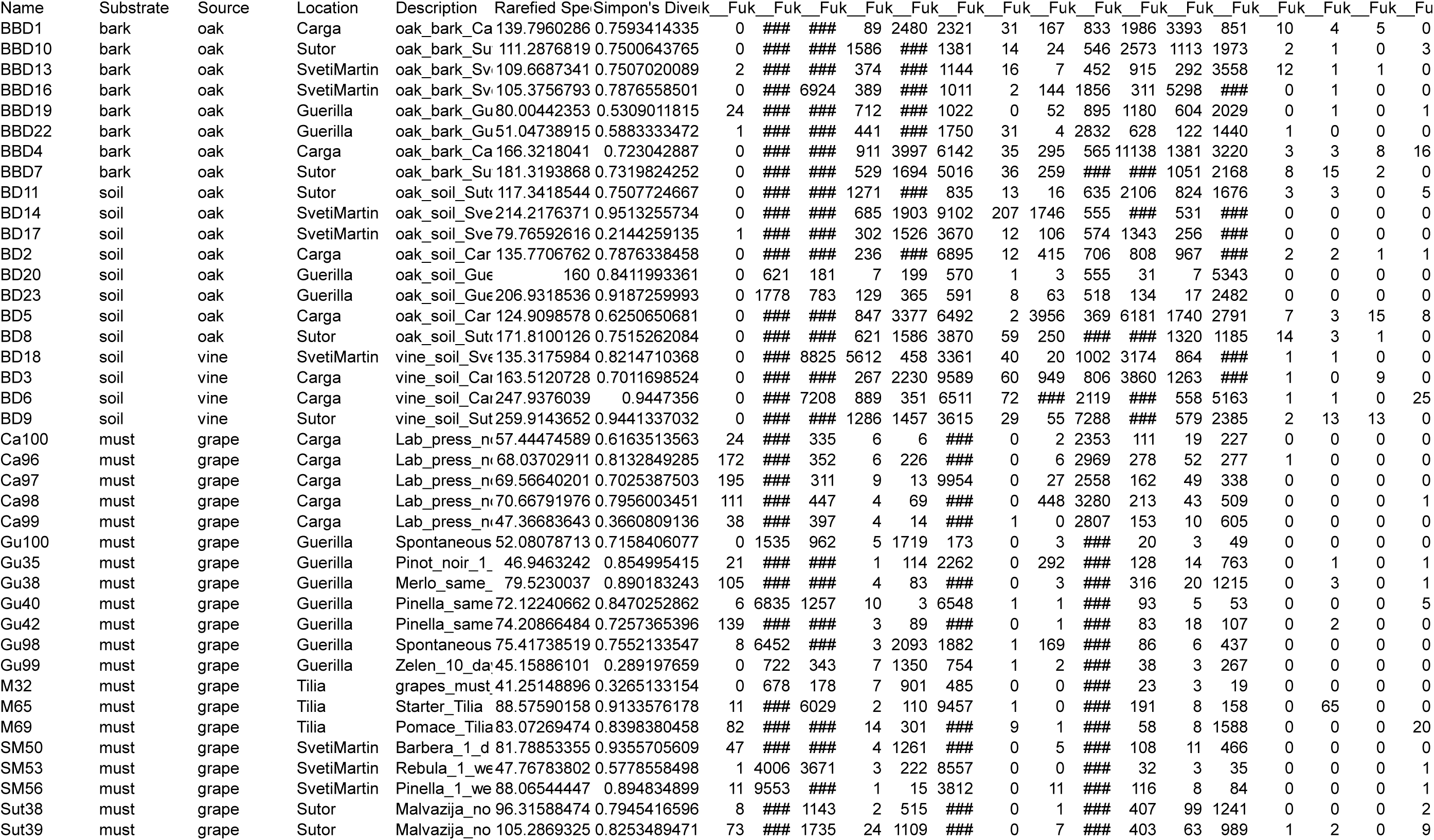
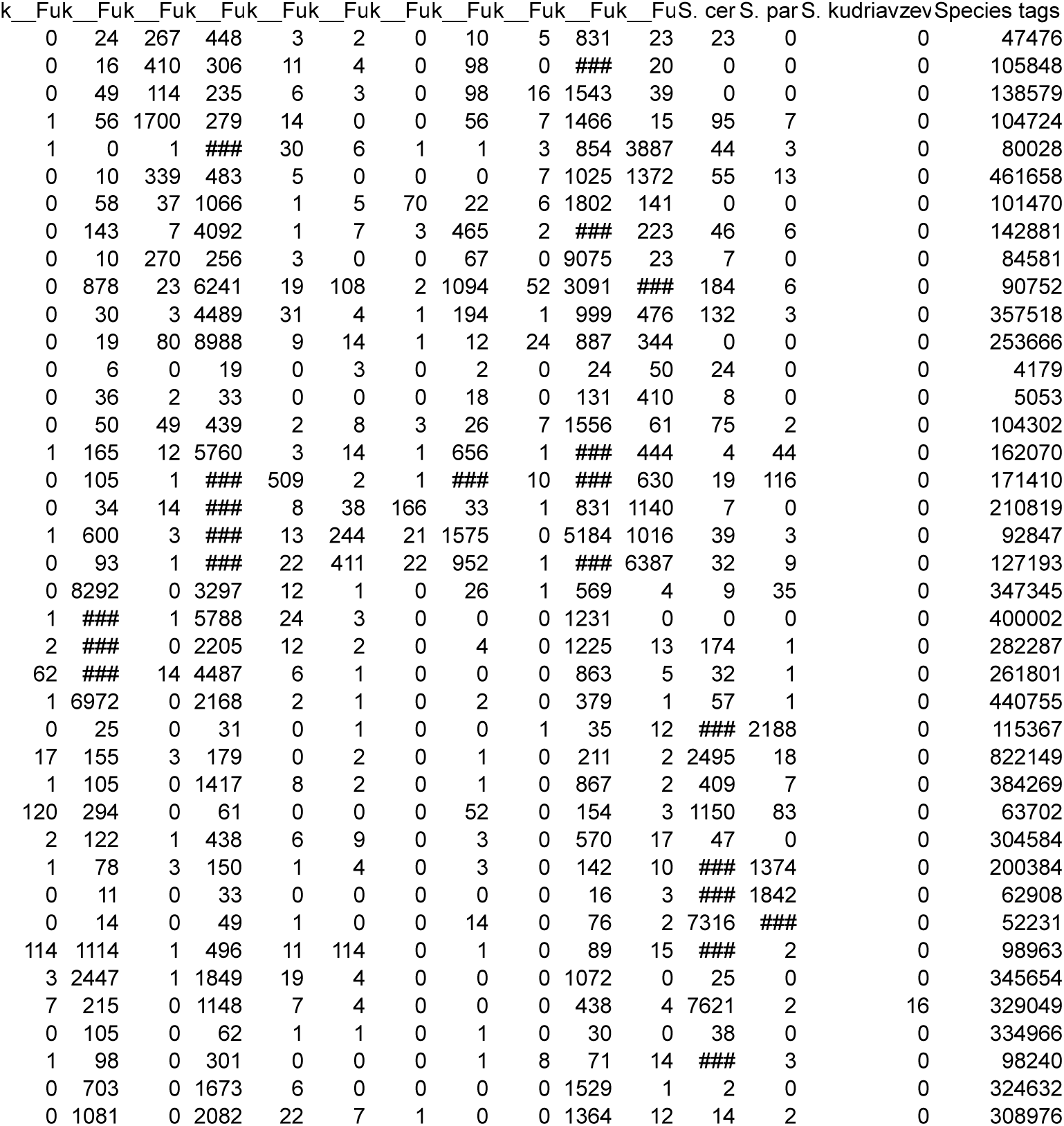
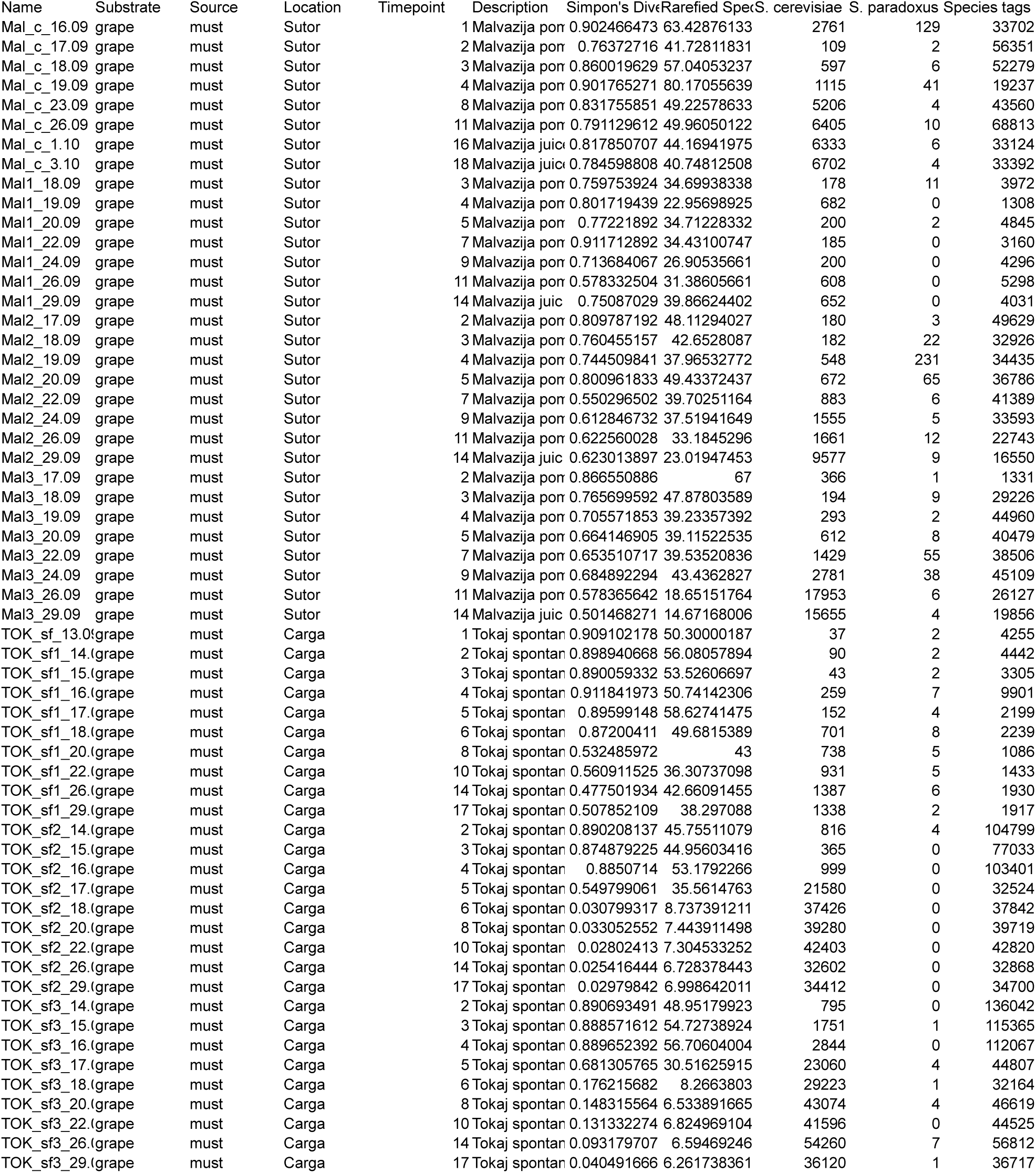
Microbiome samples, diversity and taxonomic counts.

**Table S8.** Pairwise comparisons of differences in mean levels of resistance among groups.

| Phenotype | Group | Commercial | NA Scer | NA Spar | Slovenia Scer |
| --- | --- | --- | --- | --- | --- |
| Sulfite | NA Scer | 2.00E-16 | - | - | - |
| Sulfite | NA Spar | 2.00E-16 | 0.1224 | - | - |
| Sulfite | Slovenia Scer | 0.0285 | 2.00E-16 | 2.00E-16 | - |
| Sulfite | Slovenia Spar | 2.00E-16 | 0.0027 | 1.10E-07 | 2.00E-16 |
| Copper | NA Scer | 5.40E-08 | - | - | - |
| Copper | NA Spar | 9.80E-07 | 0.54 | - | - |
| Copper | Slovenia Scer | 0.51 | 8.10E-14 | 2.00E-12 | - |
| Copper | Slovenia Spar | 3.70E-12 | 0.63 | 0.63 | 2.00E-16 |
| Low pH | NA Scer | 4.60E-09 | - | - | - |
| Low pH | NA Spar | 3.60E-15 | 0.08424 | - | - |
| Low pH | Slovenia Scer | 0.45103 | 3.90E-11 | 2.00E-16 | - |
| Low pH | Slovenia Spar | 2.00E-16 | 0.00043 | 0.19182 | 2.00E-16 |
| Ethanol | NA Scer | 0.14657 | - | - | - |
| Ethanol | NA Spar | 1.60E-05 | 0.00393 | - | - |
| Ethanol | Slovenia Scer | 0.11862 | 0.76721 | 4.20E-05 | - |
| Ethanol | Slovenia Spar | 0.00062 | 0.14657 | 0.01212 | 0.0009 |

Significance is shown by the false discovery rate based on pairwise t-tests assuming unequal variances.

**Table S9.** North American isolates of S. cerevisiae and S. paradoxus.

| Class | Site | *S. cerevisiae* | *S. paradoxus* | Sample size |
| --- | --- | --- | --- | --- |
| MO |  | 181 | 87 | 1074 |
| OR |  | 6 | 153 | 395 |
| Vineyard |  | 98 | 130 | 941 |
| Non-vineyard |  | 89 | 110 | 528 |
| Grape |  | 16 | 6 | 492 |
| Oak |  | 171 | 234 | 977 |
| 2008 |  | 88 | 205 | 947 |
| 2009 |  | 99 | 35 | 522 |
| Non-vineyard | Watrud, personal property, MO | 25 | 25 | 117 |
| Non-vineyard | Tyson Research Center, MO | 60 | 34 | 228 |
| Vineyard | Chaumette Vineyard, MO | 88 | 21 | 588 |
| Vineyard | Mount Pleasant Vineyard, MO | 8 | 7 | 140 |
| Non-vineyard | Chip Ross Park, OR | 0 | 41 | 103 |
| Non-vineyard | Bollman, personal property, OR | 4 | 10 | 79 |
| Vineyard | Tyee Vineyard, OR | 2 | 39 | 96 |
| Vineyard | Whistling Dog Vineyard, OR | 0 | 63 | 118 |

**Table S10.**
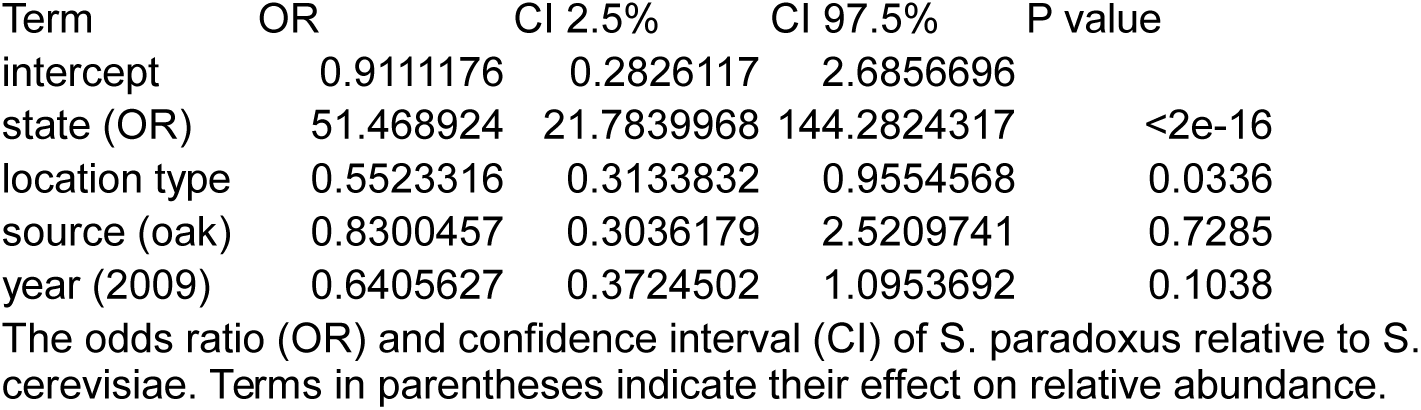
Relative abundance of S. paradoxus to S. cerevisiae in North America.

